# Variability in single-cell oxygen consumption kinetics

**DOI:** 10.1101/2023.12.06.570361

**Authors:** Ermes Botte, Yuan Cui, Chiara Magliaro, Maria Tenje, Klaus Koren, Andrea Rinaldo, Roman Stocker, Lars Behrendt, Arti Ahluwalia

## Abstract

We combined microfabricated devices with multiparameter identification algorithms to probe the variability in size-dependent oxygen consumption parameters of single human hepatic cells. We demonstrate that single cells exhibit an oxygen-dependent metabolic rate, typical of Michaelis-Menten kinetics, and that their maximal oxygen consumption is significantly lower than that of monolayers or 3D hepatic cell aggregates. Notably, we found that clusters of two or more cells competing for a limited oxygen supply reduced their maximal single-cell consumption rate, highlighting their ability to adapt to local resource availability and the presence of nearby cells. Next, we used our high-throughput approach to characterize the covariance of size and oxygen consumption within a cell population.

The results show that cooperative behaviour emerges in cell clusters, and that single-cell size and metabolism can be described by a lognormal joint probability density. Our study thus serves as a foundation to connect the metabolic activity of single human hepatocytes to their tissue-or organ-level metabolism as well as describe its size-related variability through scaling laws.

## Introduction

Biological variability (*i.e*., the fluctuation of physiological traits among individuals of the same population, also referred to as biological noise) is ubiquitous and can impact phenomena such as metabolic scaling and resilience to environmental perturbations [1, 2]. Variability is often not confined to one parameter but characterized by an interplay between multiple variables within an organism or ecosystem. For instance, it has been suggested that the covariance between individual size and metabolism conditions the ability of living organisms to react to external stimuli (*e.g*., toxins or drugs) or explain patterns in homeostatic control [1, 3, 4]. Joint variations between physiological parameters can also impact the susceptibility of organisms to diseases and their overall health [5]. Biological variability has been extensively investigated at the molecular level (transcription and expression), but less so at cellular and organismal levels. Investigating individual cells, instead of in tissues or organs, offers the opportunity to characterize variability between individuals to infer dynamics occurring at higher scales of complexity. Single cells also provide a suitable testbed to determine intrinsic size-related variability and its role in metabolic scaling, which has been highlighted as a criterion for translatability of biological parameters from micro-scale i*n vitro* systems to the *in vivo* context [1, 6, 7].

As oxygen (O_2_) is at the heart of aerobic metabolism [8, 9], several studies have measured O_2_ consumption in individual mammalian cells [10–13]. However, the accuracy, reproducibility, and throughput of O_2_ measurements at this scale remain challenging. Mammalian cells possess the ability to modulate their O_2_ consumption according to its availability, a process that is, in turn, influenced by numerous factors (*e.g*., height of culture medium and cell density) [14, 15]. However, many studies assume that cells possess a constant (zero-order) consumption rate which depends only on cell phenotype. The O_2_ consumption rate (*R*_*cell*_, in mol s^-1^) of a single cell as a function of the surrounding O_2_ concentration (*c*, in mol m^-3^) is typically represented by the Michaelis-Menten (MM) model [16–18] via two parameters: the maximal consumption rate (*sOCR*, in mol s^-1^) and the MM constant (*k*_*M*_, in mol m^-3^).

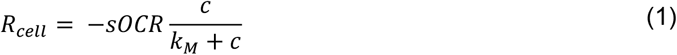

*k*_*M*_ corresponds to the concentration at which the consumption rate is half of its maximal value. At high O_2_ levels (*c* ≫ *k*_*M*_), the cellular consumption rate saturates at its maximum value (*i.e*., *R*_*cell*_ ≅ − *sOCR*). On the other hand, if *c* ≪ *k*_*M*_, the cellular uptake rate depends on the O_2_ concentration as 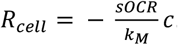. Hence, a cell with a low *k*_*M*_ value consumes O_2_ maximally even at low concentrations, whereas a cell with a high *k*_*M*_ has a low O_2_ uptake efficiency (*i.e*., O_2_ levels must be high to achieve near-maximal consumption rates).

Despite the widespread application of Eq. (1) and its relevance to the prediction and understanding of the ability of cells to adapt to environmental conditions, MM parameters for individual mammalian cells have not been measured. Here we present a systematic approach to conduct single-cell measurements of O_2_ consumption as described by the MM model. Our primary objective was to estimate the MM kinetic parameters (*i.e*., *sOCR* and *k*_*M*_) and explore their size-related variability in a human hepatic cell line, HepG2 [19]. Using custom glass microwell devices coated with luminescent O_2_-sensitive optode materials [20, 21], we isolated single cells or clusters of a few cells (from two to seven) under precisely controlled experimental conditions. Automated fluorescence microscopy was used to perform time-series imaging of the wells to extract cell sizes and O_2_ concentration profiles of individual wells. A multiparameter identification procedure [22] was applied to determine MM parameters from these profiles. Through this approach, we were able to estimate single-cell size and metabolic parameters of O_2_ as a joint probability distribution and so describe their correlated variability.

This quantitative description of single-cell O_2_ consumption allows probing the biophysical basis of cooperative metabolic dynamics, which are only detectable at higher levels of organization. Our study thus offers guidance in the design of *in vitro* or *in silico* cell-based models (*e.g*., microtissues, organoids) to improve their predictive value in biomedical applications.

## Results

Using glass microwell devices (Figure 1A) in conjunction with O_2_ sensitive optode chemistry (Figure 1B) we measured a total of 1080 microwells across five independent experiments. Of these, 227 microwells (∼ 21%) contained at least one hepatic cell whose O_2_ consumption dynamics (Figure 1D) could be recorded without interference from neighbouring wells (see SI, section SI4). The distribution of the number of cells per well (*N*_*cell*_) is reported in Figure 1E. Notably, in more than half of the cases in which cells were present in a microwell (*i.e*., 119 measurements, ∼ 11% of all investigated microwells), a single cell was probed (Figure 1C), while a maximum of seven cells per well was recognized only once.

**Figure 1.**
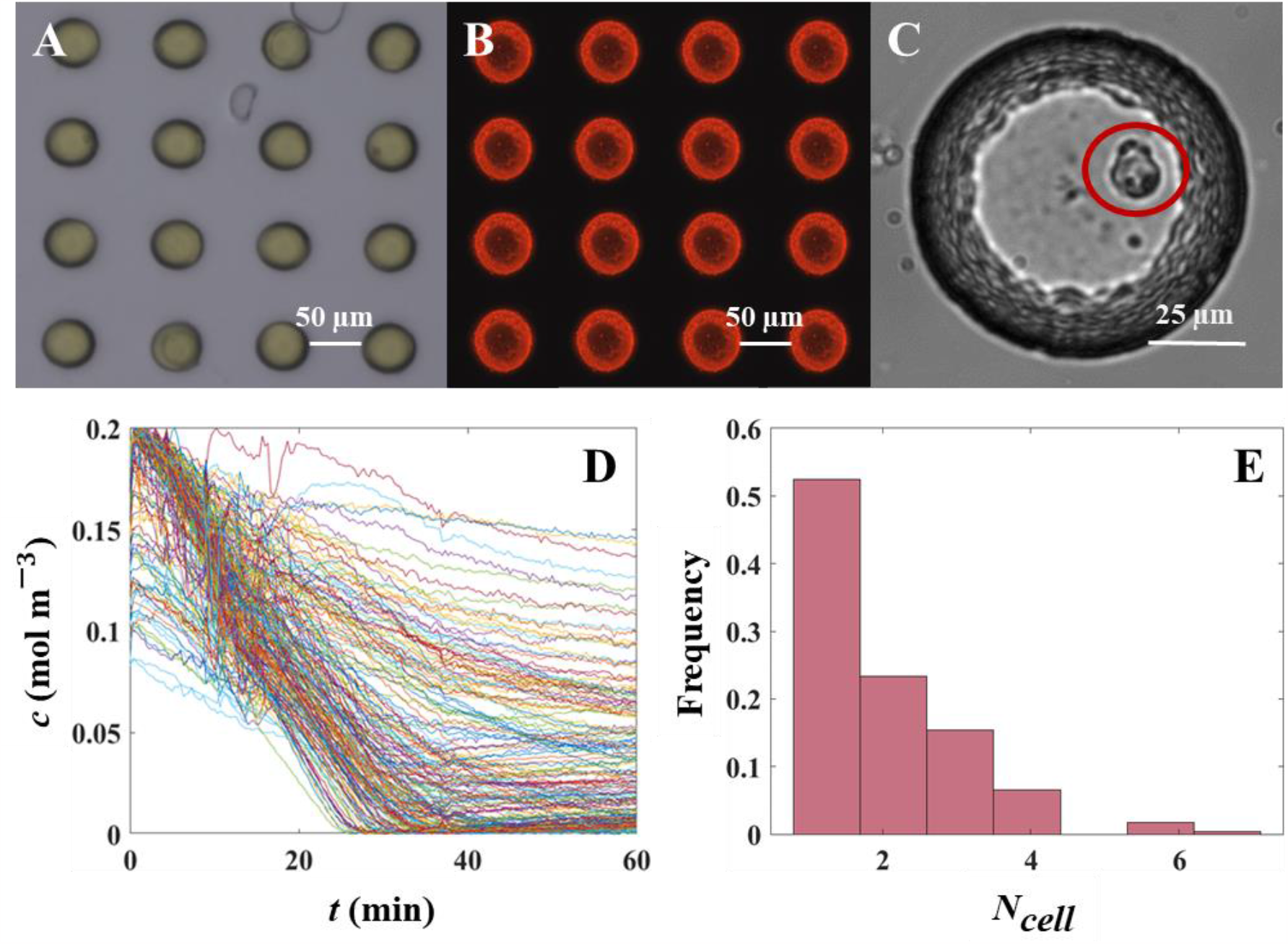
O_2_ dynamics of single hepatic cells in glass microwells. (**A**) Brightifield image of individual microwells with deposited optode chemistry (yellow) in the custom-built glass microwell array. (**B**) Corresponding luminescent response (red) upon exposure to excitation light. (**C**) Image of a single hepatic cell in a microwell with 100 μm diameter. (**D**) O_2_ concentration profiles measured in 216 microwells. (**E**) Occupation (*N*_*cell*_) frequency of cells in microwells.

### Oxygen consumption kinetics and number of cells per well

The O_2_ profiles determined from individual or multiple cells within a glass microwell were used to compute *sOCR* and *k*_*M*_ for cells within each microwell through a multiparameter identification algorithm [22]. The two MM parameters are reported both as overall probability distributions (Figure 2A, 2C) and as a function of *N*_*cell*_ (Figures 2B, 2D). While our *sOCR* values are similar to previous estimates for hepatocytes cultured in a hollow fibre bioartificial liver [23], they were significantly lower (median *sOCR* = 9.8×10^-18^ mol cell^-1^ s^-1^) than most of those reported in the literature for hepatocyte monolayers or 3D aggregates (median *sOCR* = 5.5×10^-17^ mol cell^-1^ s^-1^, Wilcoxon test, *p* < 0.0001, Figure 3A) [6, 15, 18, 22, 24–26]. Moreover, a Kruskal-Wallis test highlighted significant differences among *sOCR* values determined for microwells containing a different number of cells (*p* < 0.0001). Specifically, we found that *sOCR* decreases with increasing *N*_*cell*_ (Figure 2B, Spearman coefficient *r* = −1). This suggests that cells adjust their *sOCR* when O_2_ availability is limited because of consumption by other nearby cells.

**Figure 2.**
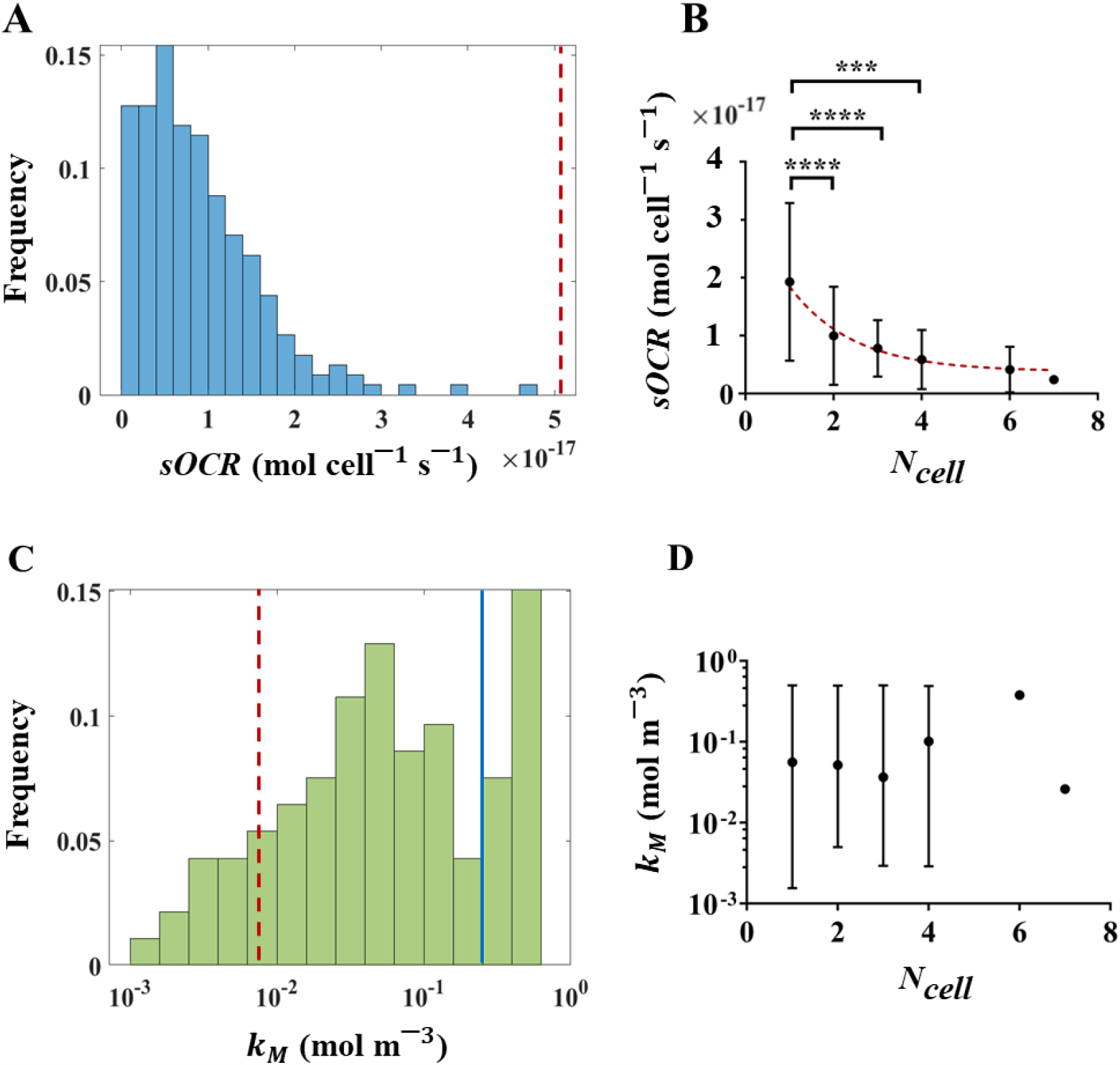
MM kinetic parameters in isolated hepatic cells determined through the multiparameter identification procedure. (**A)** Relative frequency of *sOCR* in occupied microwells. (**B)** Correlation between the number of cells per microwell and the corresponding *sOCR* value (r = −1), over all occupied microwells. The red dashed curve is a weighted fit of results (one phase exponential decay, *R*^*2*^ = 0.9797). Pairwise statistical differences computed using Dunn’s *post hoc* multiple comparisons are also reported (^***^ p < 0.0005, ^****^ p < 0.0001). (**C)** Relative frequency of *k*_*M*_ in occupied microwells (expressed in logarithmic scale). (**D)** No correlation was observed between *k*_*M*_ and the number of cells per well (r = −0.086). The vertical red dashed lines in (**A)** and (**C)** denote median literature values of *sOCR* and *k*_*M*_ [13, 16, 20–23], respectively (see Table 1 for a complete list of reported values), while the solid blue line in **(C)** indicates the O_2_ saturation level in water (*c*_0_). In (**B)** and (**D)**, data are reported as median ± range.

**Figure 3.**
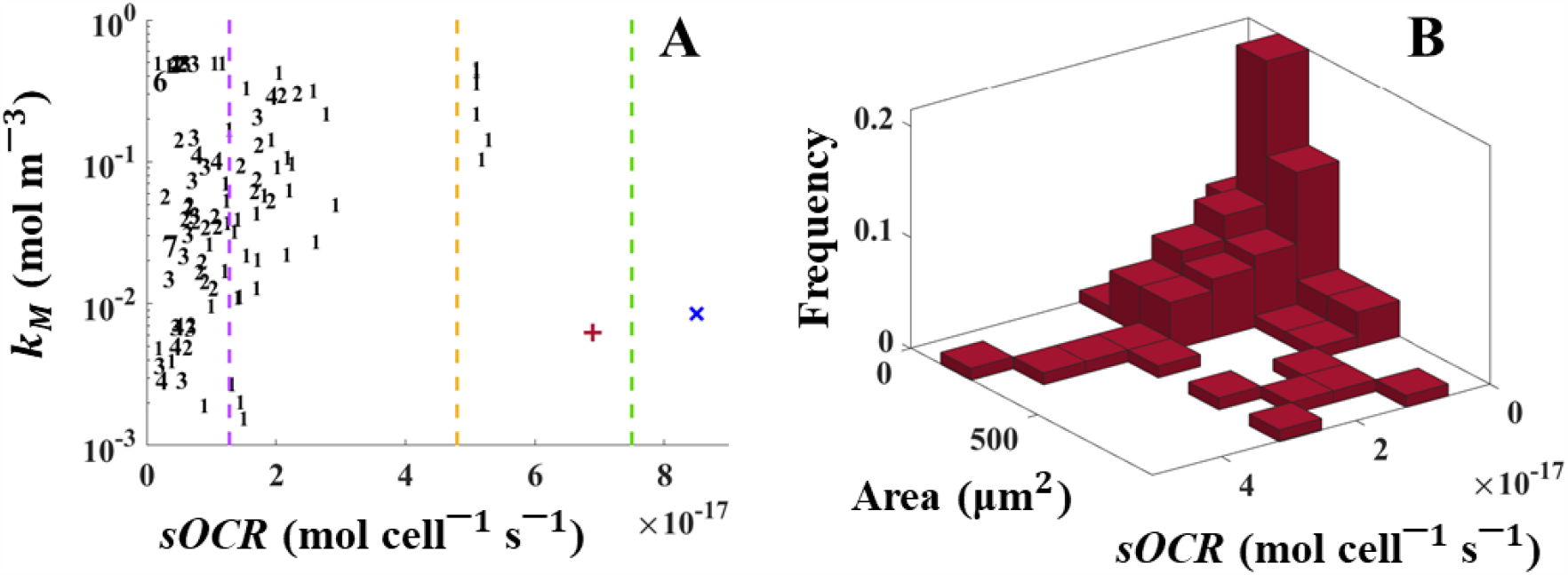
Mutual dependency of MM parameters and the size-related variability of O_2_ consumption in single human hepatic cells. **(A)** Scatter plot of measured *sOCR* and *k*_*M*_ values. The Spearman coefficient indicates that the two parameters do not exhibit any correlation (*r* = 0.1608). Each data point is denoted by a number, which corresponds to the value of *N*_*cell*_ for that data point. The red plus [15] and the blue cross [22] are median bulk values from the literature (see Table 1). Dashed lines indicate *sOCR* values from previous studies assuming zero-order kinetics for O_2_ consumption (yellow: [25], green: [26], purple: [23]). Note that *k*_*M*_ values are reported in logarithmic scale. (**B)** Joint distribution of single-cell size (*i.e*., projected area) and O_2_ metabolism (*i.e*., *sOCR*), expressed as relative frequency of occurrence.

Our measurements of *k*_*M*_ show a wide distribution, covering three orders of magnitude (Figure 2C), with a significantly higher median (5.3×10^-2^ mol m^-3^) compared to values previously reported for 2D and 3D hepatic constructs (8.5×10^-3^ mol m^-3^ – Wilcoxon test, *p* < 0.0001) [15, 22]. Approximately 36% of the measured *k*_*M*_ values are comparable to or even higher than the O_2_ saturation level in water (*c*_0_ = 0.21 mol m^-3^), suggesting that, once isolated, about a third of the cells do not approach their maximal consumption rate but instead follow first order kinetics. Due to the large variability of *k*_*M*_, neither a significant correlation with *N*_*cell*_ (Spearman coefficient *r* = −0.086) nor statistical differences among its medians for different *N*_*cell*_ values (Kruskal-Wallis test, *p* = 0.7603) were detected (Figure 2D).

**Table 1.** Values of *sOCR* and *k*_*M*_ from previous studies as bulk averages of human hepatic cell aggregates. Median values are reported for studies investigating different cell densities [15, 22]. For references which consider zero-order consumption kinetics [23–26], *k*_*M*_ values are not reported (NA).

| Aggregate type | $sOCR$ ($\times 10^{-18}$ mol cell$^{-1}$ s$^{-1}$) | $k_M$ ($\times 10^{-3}$ mol m$^{-3}$) | Reference |
| --- | --- | --- | --- |
| Isolated single cell | 9.8 | 53 | This study |
| Cell-laden hydrogel (3D) | 69 | 6.2 | [15] |
| Cell monolayer (2D) | 37 | 5.0 | [22] |
| Cell-laden spheroid (3D) | 85 | 8.5 | [22] |
| Bioartificial liver (3D) | 13 | NA | [23] |
| Cell monolayer (2D) | 55 | NA | [24] |
| Cell-laden hydrogel (3D) | 48 | NA | [25] |
| Microcarrier (3D) | 78 | NA | [26] |

To help distinguish mutual dependencies of *sOCR* and *k*_*M*_, MM parameters for all microwells investigated are presented in a scatter plot (Figure 3A). Data points referring to microwells populated by the same number of cells are clustered along the *sOCR* axis. However, no noticeable separation among groups of points corresponding to different values of *N*_*cell*_ with respect to *k*_*M*_ was observed. A correlation analysis indeed suggests that the two MM parameters do not depend on each other (Spearman coefficient *r* = 0.1608), a finding that is in contrast to previous observations for 2D and 3D aggregates of hepatic cells [22]. For comparison, bulk values for the two kinetic parameters, averaged over several cells as reported in the literature [6, 15, 18, 22–26], are also indicated in Figure 3A and listed in Table 1.

### Joint measurements of single-cell size and metabolism

Immediately after each experiment, HepG2 cells were stained with Trypan Blue, which allowed determining single-cell sizes from projected cell areas. From this data, we estimated the joint distribution of single-cell size and *sOCR* (Figure 3B). The Henze-Zirkler test (α = 0.01) demonstrated that the sample can be described by a lognormal joint probability density function (*p* = 0.0344). Further confirmation of lognormality was provided by independent optical measurements of dry mass of individual HepG2 cells – performed by means of quantitative phase imaging – which coherently displayed a marginal distribution with a lognormal shape (see SI, section SI6).

## Discussion

Here we report on the characterization of O_2_ metabolism in single human hepatic cells. Using microfabricated glass devices, coated with O_2_-sensitive optodes, one or a few cells were seeded in each microwell, enabling the precise measurement of O_2_ consumption over time as a function of the number of cells per microwell. These data were then exploited in a multiparameter identification algorithm [22] to characterize the cellular O_2_ consumption kinetics according to the MM model (Eq. (1)).

Using this integrated *in silico-in vitro* approach, we demonstrate that isolated HepG2 cells have a lower *sOCR* compared to values previously reported for 2D or 3D aggregates of the same cell type in comparable environmental conditions (Figure 2A) [15, 22, 24–26]. Additionally, our data show higher *k*_*M*_ values compared to those reported in earlier studies (Figure 2C) [15, 22]. This suggests that cells modulate their O_2_ metabolism when isolated as individuals or in clusters of a few cells. In addition, the decrease in *sOCR* with increasing number of cells per microwell, *N*_*cell*_, shows that the modulation occurs as a function of both the local O_2_ concentration and the presence of other cells (Figure 2B). This behaviour mirrors observations on hepatocytes in 2D and 3D aggregates [22] and could be interpreted as an effect of cooperation among individual cells that coexist in a microenvironment where a limited resource is shared. However, contrary to what we previously observed for hepatic cells in 2D and 3D [22], the adaptive behaviour in isolated cells appears not to affect the O_2_ uptake efficiency, since *k*_*M*_ is not dependent on *N*_*cell*_ (Figure 2D), and the two parameters are not significantly correlated (Figure 3A). It is worth noting that the identification of *k*_*M*_ is influenced by the ability of the microwell system to effectively achieve hypoxic steady-state conditions. Thus, although the optodes are highly sensitive at low O_2_ concentrations (see SI, section SI3), even slight variations of stationary O_2_ levels can influence the estimation of *k*_*M*_ and widen its distribution. Indeed, precisely characterizing *k*_*M*_ is a well-known challenge because of the sensitivity of estimated values to experimental conditions [22, 27, 28], which might ultimately mask potential trends. Nonetheless, the estimations of *sOCR* are statistically robust and not impacted by the uncertainty in *k*_*M*_, as shown in Figure 2B.

The throughput of our approach allowed us to measure the correlated variability of cell size and maximal O_2_ consumption rate (*i.e*., *sOCR*) and describe their joint frequency distribution (Figure 3B). This outcome successfully overcomes the challenge of jointly investigating individual size and metabolism, a hurdle previously encountered by our group and others [3]. Although further experiments are required to fully capture the covariance between single-cell size and O_2_ consumption [1], our study is the first size-metabolic rate distribution based on joint experimental datasets reported so far. Notably, our analysis demonstrates that the sample is extracted from a population characterized by a lognormal multivariate function. Lognormally-distributed marginal probabilities have been commonly observed for organismal sizes [29], but they have not been explicitly reported for metabolic rates.

Characterizing the covariance of single-cell size and metabolic parameters – as achieved here – is crucial to comprehensively describe the metabolic dynamics (*e.g*., O_2_ consumption) of human hepatocytes in isolation. It also paves the way to link these dynamics to the tissue or organ levels and to interpret behaviours emerging at higher scales such as cooperation or size-related scaling. A direct application of this work is the development of engineered cellular systems capable of recapitulating the heterogeneity of single-cell metabolic phenotypes. This may enhance precision medicine approaches, drug development strategies or (eco)toxicological assessments. Besides the biomedical field, the proposed methodology is also of general interest as it provides a powerful framework for systematically characterizing O_2_ metabolism and its size-related fluctuations in other single-cell scenarios, ranging from the photosynthetic activity of marine microorganisms to the evaluation of the impact of chemicals or environmental stressors on single-cell respiration.

## Materials and Methods

### Microwell devices for single-cell isolation and oxygen sensing

Single cells or small clusters of cells were isolated in a custom-built microwell array (see the schematic in Figure 4 and SI, Figure S2). The array consists of a standard borosilicate glass slide with geometrically arranged microwells (100 columns × 250 rows, *n* = 25000 microwells) fabricated via standard dry etching techniques. The microfabrication procedure for this device is summarised in the Supporting Information (SI, Figure S1), and Table S1 reports relevant technical features of the dry etching process and geometric specifications of the array customized to match the size of human hepatic cells and to minimize optical and diffusional crosstalk between microwells.

**Figure 4.**
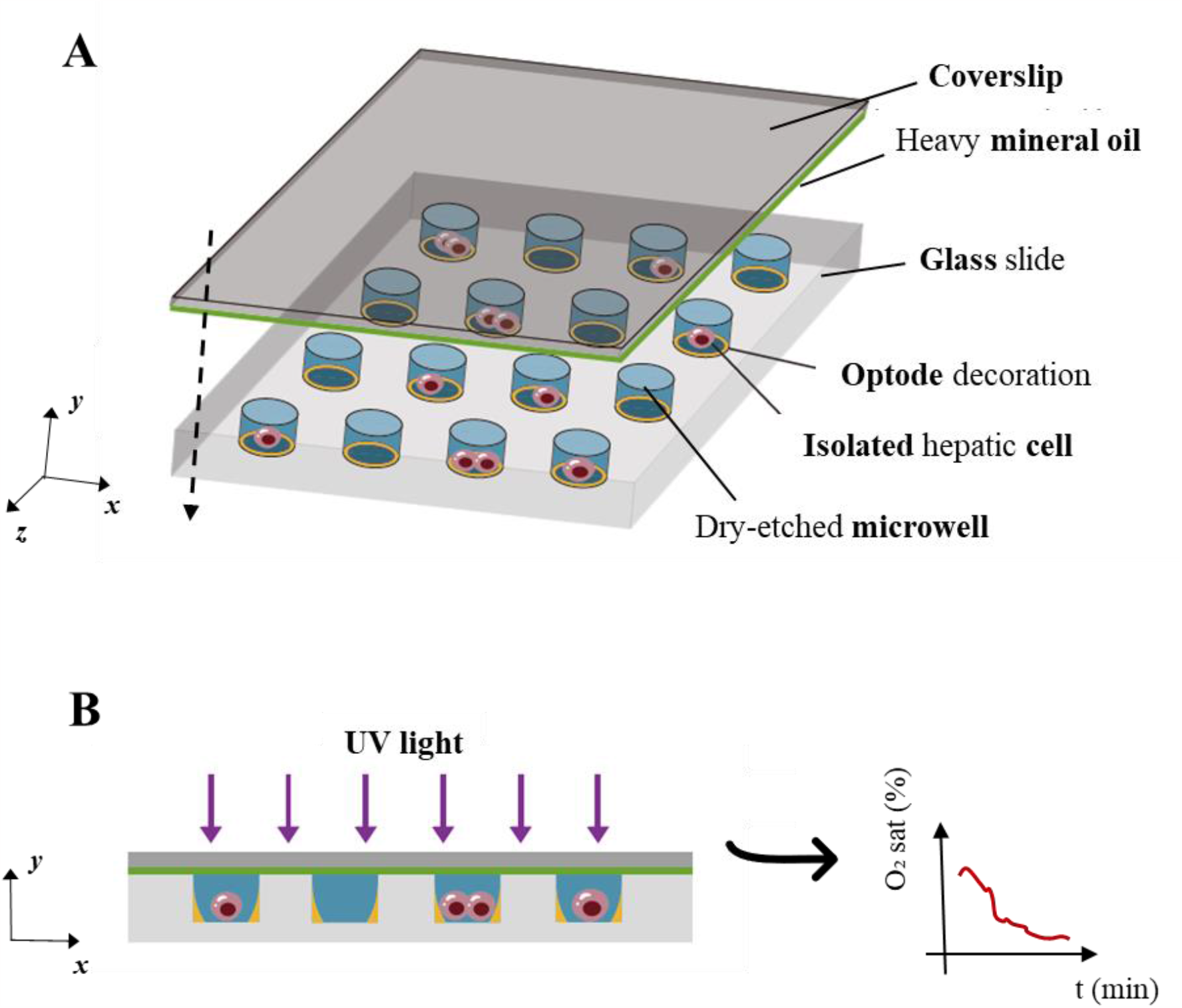
Schematic of the assembled glass microfluidic device. **(A)** Microwells coated with optode chemistry are seeded with hepatic cells suspended in culture medium, and subsequently the device is overlaid with a glass coverslip coated in heavy mineral oil. This oil interface reduces lateral (*i.e*., inter-well) O_2_ diffusion so that the microwells effectively reach hypoxic conditions. **(B)** Once sealed, the system is exposed to short pulses of UV light and the resulting luminescence emission from optodes is detected over time to derive O_2_ concentrations across microwells. Inset: a typical O_2_ concentration profile obtainable by monitoring optode response in an individual microwell where at least one cell is settled.

### Optode sensor composition, deposition and calibration

#### Composition

Quenching-based luminescent optode materials were chosen for O_2_ sensing as they are characterized by a high spatial resolution and short response time, as required for single-cell O_2_ consumption rate measurements [30, 31]. The deposited optode material was composed of platinum(II)-5,10,15,20-tetrakis-(2,3,4,5,6–pentafluorophenyl)-porphyrin (PtTFPP), polystyrene (PS) and MACROLEX^®^ yellow 10GN (MY) dissolved in toluene. Here, PtTFPP is the O_2_-sensitive dye whereas MY acts as a reference (*i.e*., O_2_-insensitive) dye. Both dyes are excited using UV light (λ = 396 nm) and emit red luminescence (PtTFPP, λ ≥ 650 nm) or green luminescence (MY, λ = 507 nm) in an O_2_ concentration-dependent manner [31] or at a constant intensity, respectively. The ratio of red-to-green (R/G) emission intensities thus represents a robust O_2_-dependent signal, with minimal noise from environmental artefacts (*e.g*., sensor bleaching) and, with appropriate calibration, allows for relating to O_2_ contents within individual microwells.

#### Deposition

To deposit the optode material, a 10% w/v solution of PS in toluene containing 0.15 g/L of both MY and PtTFPP was spread on the glass microwell array via a pipette. The homogeneity of the optode coating was assessed by profilometry (see SI1 for details), which revealed that it had a nominal thickness of 5 μm and was most uniform in the central region of the array. Therefore, we prioritized this central area (containing at least 200 microwells) for seeding cells and investigating their O_2_ consumption. As the deposited optode material is hydrophobic, the array was briefly treated with O_2_ plasma (Zepto, Diener electronic, Ebhausen, Germany) for 10 s at 0.4 mbar and 50% intensity to promote filling of microwells with culture medium and assist in cell adhesion. This plasma treatment did not affect the response of optode materials to O_2_ (see SI2 for details). To avoid undesired background fluorescence, the optode material deposited outside of the microwells was removed using a scalpel. This process resulted in two regions with different wettability, *i.e*., (i) hydrophilic microwells containing a layer of optode material (yellow regions in the inset of Figure 1A) surrounded by (ii) hydrophobic glass (gray regions in the inset of Figure 1A).

#### Calibration

Following optode deposition, a calibration curve was constructed by averaging data acquired from 216 central microwells (see SI1 for details). To perform calibration measurements, the optode-coated array was placed into a gas-impermeable chamber with a transparent window for image acquisition. The chamber was placed in a microscopy incubator (Okolab srl, Pozzuoli, Italy) maintained at 37°C, to avoid temperature fluctuations, which might influence optode responses (Figure S2). Optode emissions were recorded via a fully automated fluorescence microscope (Nikon Ti2-*E*, Nikon, Tokyo, Japan) equipped with an RGB camera (DFK 33UX264 colour industrial camera, The Imaging Source) and a LED excitation light (Spectra X, Lumencor, OR, USA).

Calibration was performed by introducing gases at known levels of O_2_ saturation. Specifically, the R/G in totally anoxic conditions (*R*/*G*_0_) was measured by exposing the device first to pure nitrogen (N_2_) and thereafter to compressed air (*i.e*., 21% O_2_) to obtain 100% air saturation (*R*/*G*_100_), corresponding to an equilibrium concentration in aqueous media equal to *c*_0_. Images of single microwells were acquired with a 40× objective. All other imaging parameters were set as listed in Table 1 for monitoring cellular O_2_ consumption. The measurement was used to derive the Stern-Volmer (SV) quenching constant [31] (*k*_*SV*_, expressed in 1/% air sat.) of the array as follows.

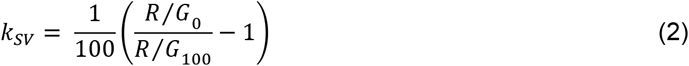

Calibration curves were also estimated in liquid phase (*i.e*., cell culture medium) and compared to those obtained for corresponding microwell arrays using gases (see SI2 for details). No significant differences emerged between liquid phase and gas phase calibrations, and hence we routinely relied on the more convenient gas calibration prior to each experiment. Further, we ensured that optode bleaching was negligible in the timeframe of our single-cell O_2_ consumption tests (see SI3 for details). Finally, experiments were conducted to determine the extent of optical crosstalk between microwells and to define inter-well distance and cell occupancy (see SI4 for details).

### Single-cell oxygen measurements

#### Cell preparation

Human hepatic cells from hepatocellular carcinoma (HepG2 cells, ATCC, Manassas, Virginia, USA) were maintained in T25 flasks (Sarstedt, Numbrecht, Germany) under standard conditions (37 °C, 95% humidity, 5% CO_2_) and supplied with fresh Dulbecco’s Modified Eagle Medium (DMEM, Sigma-Aldrich, St Louis, Missouri, USA) every 3 days. Before experiments, cells were detached with trypsin-EDTA (Lonza, Basel, Switzerland).

After optode calibration, the central area of the microwell array was overlaid with a thick poly-dimethyl-siloxane (PDMS) frame into which 500 μL of cell suspension in DMEM was pipetted (Figure 1B). This allowed control of the seeding density and ensured that cells were confined to the region where the optode coating was determined to be homogeneous. Experiments were performed with different seeding densities, ranging from 2×10^3^ cells mL^-1^ up to 10^6^ cells mL^-1^. The former resulted in an optimal trade-off between an acceptable number of microwells containing single cells and minimal lateral O_2_ diffusion between neighbouring microwells. Following cell seeding, the device was incubated overnight to allow cell adhesion to the microwells (inset in Figure 1B).

#### Monitoring single-cell consumption

Just before the single-cell O_2_ consumption experiments, the culture medium within the PDMS frame was changed and 60 μL of fresh DMEM were dispensed into the microwells. Then, the PDMS frame was removed and microwells seeded with cells were covered with a coverslip, which was affixed to the array via both paperclips and magnets (Figure 1A). A thin coating of heavy mineral oil (Sigma-Aldrich) was applied to the underside of the coverslip to isolate the microwells from environmental O_2_ (Figure 4) and to define the initial and boundary conditions of the system. Mineral oil has a significantly lower O_2_ diffusion coefficient than aqueous media [32] and effectively seals off the array from ambient O_2_, allowing a hypoxic steady state to be reached – a necessary condition to properly characterize the MM consumption kinetics, particularly *k*_*M*_ (see subsection *Modelling single-cell consumption)*. The ratio of mineral oil to cell culture media was optimized to ensure maximal phase separation and minimize lateral (*i.e*., inter-well) O_2_ diffusion and, at the same time, guarantee that cells are exposed to the medium phase within microwells (see SI5 for details). Image acquisition was started immediately after device assembly to ensure rapid monitoring of O_2_ dynamics (*i.e*., from the initial condition of maximum O_2_ in the medium to the achievement of stationary hypoxia, defined as *c* ≤ 0.04 mol m^-3^ [33]). Measurement duration and sampling frequency were set according to instrument limits and modelling considerations (see subsection *Modelling single-cell consumption*). The O_2_ dynamics associated with cell consumption were measured by repeatedly scanning the central area containing cells, using large area scanning mode. The large image acquisition parameters are listed in Table 2.

**Table 2.** Experimental parameters for O_2_ sensing, using a Nikon Ti2-*E* automated fluorescence microscopy system.

| Parameter | Numerical value |
| --- | --- |
| Magnification | 10× |
| Large image size (n×m) | 6×6 |
| Time scale ($t_{\text{end}}$) | 1 h |
| Time resolution ($\Delta t$) | 20 s |
| Excitation light wavelength | 396 nm |
| Excitation light intensity | 20% of maximum intensity* |
| Excitation duration | 50 ms |

\*To achieve optimal saturation of the sensing dye, the intensity of the excitation light was coherently adjusted for each round of calibration and subsequent experiment.

Following each experiment, 30 μL of trypan blue dye (0.4% w/v solution, Sigma-Aldrich) was gently pipetted through the gap between the microwell array and the coverslip. Trypan blue was left to diffuse over the array for about 10 min, then a brightfield large image was acquired setting the same scanning pathway as used during experiments. This procedure enabled identifying *N*_*cell*_ in each microwell and estimating cell size (*i.e*., the projected area) using ImageJ [34].

#### Determination of concentration profiles

Time series of large images were processed exploiting algorithms purposely developed in Matlab (R2021b, MathWorks, Massachusetts, USA). Briefly, each RGB image was segmented by means of customized thresholding to distinguish PtTFPP-decorated microwells from the background. This allowed for the computation of the pixel-by-pixel R/G. Finally, for each microwell, R/G profiles were determined over time by averaging over pixels belonging to the same microwell. This allowed for conversion of R/G signals into O_2_ concentrations using previously established calibration curves.

#### Modelling single-cell consumption

Leveraging analytical considerations and bearing in mind that neither cell location nor the number of cells per well (*N*_*cell*_) can be determined *a priori*, we established the duration (*t*_*end*_) and sampling frequency (*f*) of measurements based on the experimental setup. From a modelling point of view, each microwell is a region of the space governed by the reaction-diffusion equation (Eq. (3)):

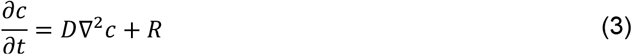

where *c* = *c*(*x, y, z, t*) [mol m^-3^] is the O_2_ concentration field in the microwell, *D* [m^2^ s^-1^] is the diffusion constant of O_2_ and *R* = *R*(*x, y, z, t*) [mol m^-3^ s^-1^] is the O_2_ production/consumption rate per unit volume. In the case of consumption, *R* is negative and can be described as a function of the single-cell MM consumption rate (*R*_*cell*_) defined in Eq. (1). Thus:

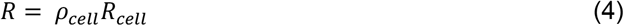

where *ρ*_*cell*_ [cell m^-3^] is the cell density in the microwell volume. Given that cell and microwell sizes are comparable, and assuming that O_2_ consumption is uniform in the cell volume – and hence in the well domain – the cell density can be expressed as 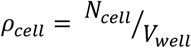 (with *V*_*well*_ indicating the microwell volume, Table S1). As the silicone oil layer is considered impermeable to O_2_, there is no flux at the oil/air boundary, likewise at the microwell walls or – given the homogeneity of consumption in the volume – within the domain. These assumptions imply that the O_2_ concentration field in the microwell depends only on time (*c*(*x, y, z, t*) = *c*(*t*)), and the governing equation can be simplified as 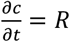. In these conditions, a suitable duration for monitoring O_2_ consumption to hypoxia within each microwell was estimated based on the characteristic reaction time, *τ*_*r*_:

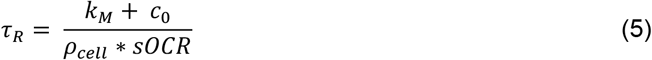

where conditions of O_2_ saturation (*i.e*., *c* = *c*_0_, see Table 3) and *N*_*cell*_ = 1 were assumed. These are cautious choices leading to the longest *τ*_*R*_ possible for the system, guaranteeing that the consumption dynamics are fully captured. Typical literature values were then used for *sOCR* and *k*_*M*_ [6, 18, 23, 24], giving *τ*_*R*_ = 678.4 s. Thus, experiments were set with *t*_*end*_ ≥ 5*τ*_*R*_ and *f* ≥ 10/*τ*_*R*_.

**Table 3.** Initial condition and range of parametric sweep implemented for simulating O_2_ dynamics within a microwell where *N*_*cell*_ cells have settled.

| Parameter | Numerical value | Description |
| --- | --- | --- |
| $c_0$ | 0.2 mol m<sup>-3</sup> ([18]) | Initial condition of $O_2$ saturated-culture medium |
| $sOCR$ | [10<sup>-18</sup>; 10<sup>-16</sup>] mol cell<sup>-1</sup> s<sup>-1</sup><br>(deduced from [18, 35]) | Initial range of parametric sweep |
| $k_M$ | [10<sup>-3</sup>; 10<sup>-1</sup>] mol m<sup>-3</sup> (deduced from [17, 18, 35]) | Initial range of parametric sweep |

### Kinetic parameter identification

Experimentally measured O_2_ concentration profiles constituted the input datasets for the multiparameter identification algorithm reported in [22]. Briefly, values of *sOCR* and *k*_*M*_ were estimated comparing the O_2_ dynamics measured in each microwell containing cells to those predicted *in silico* by modelling the system according to Eqs. (3) and (4). A model governed by the dimensionless form of Eq. (3) was iteratively solved for each specific microwell, taking *N*_*cell*_ from the trypan blue-stained image and parameters listed in Table 2 into account.

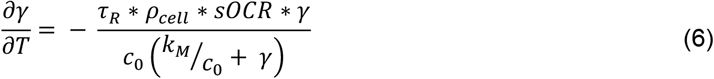

In Eq. (6), *γ* = *c*/*c*_0_ and *T* = *t*/*τ*_*R*_ are the non-dimensional concentration and time, respectively. Considering Eq. (5), the dimensionless equation implemented for simulating O_2_ consumption in the well is the following:

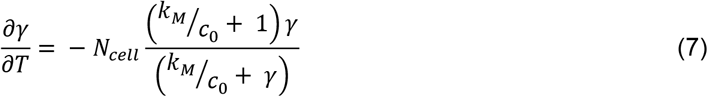

The multiparameter identification algorithm described in [22] was used to estimate the MM parameters through Eq. (7). Although the equation does not explicitly depend on *sOCR*, the latter determines the time scale of the solution, given the definition provided for the dimensionless time *T*.

### Statistical analysis and software

Numerical values of consumption parameters identified for microwells as a function of *N*_*cell*_ were compared by means of a non-parametric Kruskal-Wallis test and pairwise *post hoc* Dunn’s multiple comparisons. Non-linear correlation (*i.e*., non-parametric Spearman coefficient) of *sOCR* and *k*_*M*_ with respect to *N*_*cell*_ as well as with each other was also tested. Overall statistical differences between the MM parameters estimated here and typical literature values were assessed for both *sOCR* and *k*_*M*_ using a non-parametric Wilcoxon signed-rank test. Furthermore, the joint distribution of single-cell size and *sOCR* was tested for both normality and lognormality via the Henze-Zirkler multivariate normality test performed on the original and log-transformed dataset, respectively. Image processing, simulation of O_2_ transport and consumption and multiparameter identification were implemented in Matlab (R2021b), while GraphPad Prism (version 7, GraphPad Software, California USA) was used to perform all statistical analyses.

## Acknowledgments

The work leading to these results has received funding from the Swiss National Science Foundation, through Project SINERGIA 2019 572 CRSII5_186422/1.

We acknowledge Myfab Uppsala for providing facilities and experimental support required to fabricate the microwells. Myfab is funded by the Swedish Research Council (2019 - 00207) as a national research infrastructure.

LB was supported by grants from the Swedish Research Council (2019-04401), the Science for Life Laboratory and the Carl Trygger Foundation (CTS 20:214).

EB also acknowledges the European Union for his post-doctoral fellowship, funded by the Next Generation EU project CN00000013 ‘Centro Nazionale 1’ HPC, Big Data and Quantum Computing (CN1, PNRR, Spoke 6: Multiscale modelling and engineering applications).

## Author Contributions

AA and LB conceived the study and supervised the research and manuscript editing. EB and YC conducted experiments and performed image processing and data analysis. All authors contributed to data interpretation. EB drafted the manuscript, and all authors reviewed it as well as agreed to its submission.

