## Supplementary Information for "Variability in single-cell oxygen consumption kinetics"

### **Supporting Information for**
### Variability in single-cell oxygen consumption kinetics

Ermes Botte[a,b,1], Yuan Cui[c,d,1], Chiara Magliaro[a,b], Maria Tenje[d], Klaus Koren[e], Andrea Rinaldo[f,g], Roman Stocker[h], Lars Behrendt[c,2,*] and Arti Ahluwalia[a,b,2,*]

[a] Research Centre "E. Piaggio", University of Pisa, Italy
[b] Department of Information Engineering, University of Pisa, Italy
[c] Department of Organismal Biology, Science for Life Laboratory, Uppsala University, Sweden
[d] Department of Material Science and Engineering, Science for Life Laboratory, Uppsala University, Sweden
[e] Aarhus University Centre for Water Technology, Department of Biology, Aarhus University, 8000 Aarhus, Denmark
[f] Laboratory of Ecohydrology, École Polytechnique Fédérale de Lausanne
[g] Department of Civil, Environmental and Architectural Engineering, University of Padova
[h] Institute for Environmental Engineering, ETH Zurich, Zurich, Switzerland

[1] shared first authorship
[2] shared senior authorship

[*] corresponding authors:

**Supporting Information text**

SI1 – Characterization of the homogeneity of optode deposition

The bottom of the microwells was probed using profilometry [1] before and after application of the optode film to quantify its homogeneity and evaluate the reproducibility of the deposition process. A standard contact profilometer (Dektak 150 surface profiler, Bruker Nano Analytics, Billerica, Massachusetts, USA) based on the deflection of a diamond-tipped cantilever was used to determine the spatial distribution of the effective thickness as point-by-point height differences. The assessment was repeated for 21 microwells in six independently decorated devices, resulting in an overall effective thickness of 5 ± 3 µm (expressed as mean ± standard deviation for all assessed microwells). Most of variability in thickness was observed around the edges of the slide where optode accumulation predominates. For this reason, calibration and single-cell respiration measurements were conducted as far as possible from the edge of the array and each decorated microwell device was calibrated independently.

SI2 – Impact of plasma treatment and phase of calibration medium on optode response

As plasma treatment (see the main text, sub-section *Deposition*) could affect the sensitivity of deposited optode chemistry, we assessed this effect empirically. Calibration curves for 20 empty microwells were conducted according to the Stern-Volmer (SV) model [2], before and after treatment. The calibration was performed both in gas as well as liquid, to determine whether optode materials would behave differently. Calibration under gas is described in the main text (see sub-section *Calibration*) and a similar two-point measurement protocol was set up for calibration under liquid phase. Here, culture medium (DMEM, Sigma-Aldrich) was dispensed onto the microwell array, which was subsequently closed with the silicone oil-coated coverslip. Specifically, to determine the first point of the calibration curve, 60 µL of 100% air-saturated DMEM – corresponding to an $O_2$ concentration of $c_0$ (see Table 2 in the main text) – was obtained following the standard procedure suggested by producers of commercial quenching-based $O_2$ sensors [3]. The second calibration point – corresponding to anoxia – was achieved by preparing a DMEM calibration solution containing an $O_2$ scavenger (NaHSO$_3$, Sigma-Aldrich) at a concentration of 1% w/v [4, 5]. For all calibrations, images of single microwells were acquired using a 40× objective (see Table 1 in the main text for imaging parameters). Images were processed in Matlab (release 2021b), and $k_{SV}$ determined for each microwell through Eq. (5) reported in the main text.

Numerical values computed for parameters characterizing the SV curve (*i.e.*, $R/G_0$ and $k_{SV}$) in each configuration were statistically analysed in GraphPad Prism (version 7) using a 2-way ANOVA and *post hoc* Turkey's multiple comparisons.

As shown in Figure S3, the calibration curves do not differ significantly (α = 0.05), which confirms that neither the plasma treatment nor the phase (*i.e.*, air or water) of the calibration medium affect sensor performance. Moreover, this result indicates that sensor calibration can be more conveniently carried out in the gas rather than the aqueous phase.

SI3 – Characterization of optode bleaching

$O_2$-sensitive optode materials like PtTFPP are reported to photo-bleach or produce singlet $O_2$ when exposed to excitation light for long times [6, 7]. To compensate for potential artefacts arising from this phenomenon, optode responses were characterized over time, at multiple constant levels of air saturation (% air sat). Briefly, experimental measurements were performed in the same gas-impermeable chamber used for calibration purposes (see subsection *Calibration* in the main text) and setting five different air saturation levels by combining compressed air and $N_2$ through a gas mixer set up (Red-y smart series mass flow meters, Voegtlin GmbH, Muttenz, Switzerland). This system enabled modulation of the partial pressures of the mixture components by tuning the flow rate from their sources. The stability of air saturation within the sealed chamber was verified by monitoring it with a calibrated optical microsensor (OXR250, Pyroscience GmbH, Aachen, Germany). Optode responses were measured for 20 microwells over time, employing the same experimental parameters (Table 1, main text). The results of this optode bleaching experiment are summarized in Figure S4. Remarkably, sensor bleaching causes a measurement drift limited to 1% air sat/h across $O_2$ levels, increasing up to 2% air sat/h only for 100% air saturation (Figure S4A). This characterization also highlights the non-linearity of optode-based $O_2$ sensing, with a well-known higher sensitivity at low $O_2$ concentrations (inset in Figure S4B). This super-linear behaviour makes the current sensor chemistry particularly suitable for biomedical applications. The accurate measurement of $O_2$ concentration in hypoxic microenvironments is crucial, for instance, in the development of new cancer treatments [8–10] or for probing the cell response in $O_2$-depleted regions potentially arising in tissue models *in vitro* [11–14].

SI4 – Characterization of optical crosstalk

Another aspect that could potentially interfere with the accuracy of $O_2$ concentration measurements is the optical crosstalk between luminescent optode materials deposited within adjacent microwells in the array. This may be caused by the radially directional spreading of light emitted by both the red $O_2$ sensitive dye (PtTFPP) and the green reference dye (MY) as well as its scattering due to the surface roughness of microwells. To assess this contribution to the optical signal detected within microwells, we acquired images from 20 individual microwells exposed to excitation light (see Table 1 in the main text). Image acquisition was performed with a 40× objective under fully anoxic conditions, in order to measure the spatial characteristics of PtTFPP emission at maximum intensity, while all remaining parameters were identical to values provided in Table 1 in the main text.

The images were processed in Matlab (release R2021b). Given the isotropic nature of light scattering, polar symmetry was assumed, and the spatial decay of luminescence intensity (*I*) was modelled as a function of the radial coordinate (*r*) according to the following relationship:

$$I(r) = I(0)e^{-r/\delta} \tag{S1}$$

where $I(0)$ is the light intensity measured at the edge of the microwell (corresponding to the arbitrarily chosen origin of the radial axis, see Figure S5), and $\delta$ (µm) denotes the characteristic radial distance, *i.e.*, the distance from the microwell edge at which light intensity is damped by a factor equal to $1/e$. Eq. (S1) was used to fit experimental profiles (Figure S5) and thus determine $\delta$ for both the red and green dye. More in detail, in both cases, $\delta$ was computed as the weighted average of those obtained by fitting the optical signal from each microwell.

$$\delta = \sum_{i=1}^{20} w_i \delta_i \tag{S2}$$

In Eq. (S2), $\delta_i$ is the characteristic radial distance estimated for the *i*-th microwell, and $w_i$ represents the associated weight, defined depending on the goodness of fit as follows:

$$w_i = \begin{cases} \sqrt{{R^2}_i / \sum_{i=1}^{20} {R^2}_i}, & {R^2}_i \geq 0.75 \\ 0, & {R^2}_i < 0.75 \end{cases} \tag{S3}$$

where ${R^2}_i$ is the determination coefficient of the exponential fitting for the *i*-th microwell.

According to this rationale, we estimated characteristic distances $\delta_R$ = 51.7 µm and $\delta_G$ = 46.4 µm for PtTFPP and MY, respectively. Hence, if *d* denotes the inter-well distance (or, equivalently, the microwell diameter), considering design specifications reported in Table S1 and applying Eq. (S1) in the worst case (*i.e.*, PtTFPP under 0% $O_2$), at the edge of a neighboring well (*i.e.*, at $r = d$) the damping of luminescence intensity is given by $\frac{I(d)}{I(0)} \cong \frac{1}{e} \cong \frac{1}{3}$ for 50 µm diameter microwell arrays (*i.e.*, having *d* = 50 µm) and $\frac{I(d)}{I(0)} \cong \frac{1}{e^2} \cong \frac{1}{10}$ for 100 µm diameter ones (*i.e.*, having *d* = 100 µm). Based on these rough calculations, only adjacent wells with a diameter of 100 µm can be reliably assumed as optically decoupled (*i.e.*, $I(d) \ll I(0)$). Hence, in this study, devices with a microwell diameter of 100 µm were employed to measure the $O_2$ consumption of isolated cells.

SI5 – Optimization of oil-medium phase separation

Finally, the optimal ratio between culture medium and silicone oil used during experiments was evaluated. To this end, Nile Red (a lipophilic fluorescent dye with excitation wavelength 552 nm and emission wavelength 636 nm) was added to silicone oil to obtain a 0.001% w/v solution, and images of the device closed with the coverslip were acquired to assess the homogeneity of the oil layer, while varying the quantities of both components.

The best trade-off between culture medium and silicone oil (Figure S6A) was obtained when using 60 µL of fresh DMEM dispensed onto the array and 60 µL of silicone oil coating the coverslip. The plasma treatment - performed *a priori* – facilitated the filling of microwells with the aqueous phase (*i.e.*, DMEM), while silicone oil tended to coat the inter-well spaces and overlay the culture medium (Figure S6A). The balance between the two is crucial. If the amount of silicone oil is too low, part of the array was left uncovered (Figure S6C), reducing the sealing effect. On the other hand, when too much oil was applied, it resulted in the displacement of DMEM (and cells, if any are present) out of the microwells (Figure S6B).

SI6 – <u>Dry mass measurements of human hepatic single cells</u>

Single-cell dry masses of HepG2 cells were estimated using quantitative phase imaging (QPI). QPI leverages on physical principles of phase contrast microscopy, which exploits local phase shifts (or, equivalently, time delays) on reflected light waves. The shifts are induced by a specimen in the light path, and can hence be used for reconstructing its geometry, without application of any optical labels [15]. In fact, biological samples such as single cells or tissues do not damp the amplitude of electromagnetic fields in the visible range but are able to differentially shift their phase according to point-by-point variability in optical density, producing a phase map which can be compared to a proper reference for the image formation. In this study, a QPI camera system (SID4 sC8, PHASICS SA, Saint-Aubin, France) was integrated onto the same automated microscope used for characterizing single-cell $O_2$ consumption. To measure single-cell dry masses, 250 µL of a HepG2 cell suspension (at an arbitrary density) were dispensed onto a glass slide, closed with a coverslip and sealed with Scotch tape to avoid medium losses. Images were acquired for two different samples, either i) immediately after seeding or ii) after allowing cells to adhere to the glass during an overnight incubation. To obtain a reference for background subtraction, an area with no cells was first imaged, and subsequently individual cells were manually selected to assess their volume (and, thus, dry mass) via a proprietary software (sid4Bio, PHASICS SA) associated with the camera. All images were acquired using a dedicated 60× objective (Plan Apo – 60×/0.95, Nikon).

Collected datasets of HepG2 cell dry masses were processed using Matlab (release 2021b). Firstly, the Anderson-Darling (AD) test was performed on dry mass distributions of both suspended and adhered HepG2 cells (Figure S7A), demonstrating that the two distributions can be described by the same unspecified functional form ($\alpha = 0.01$), corroborating the robustness of the approach with respect to isovolumetric changes of shape. This is coherent with cell incompressibility. In fact, mammalian cells are known to spread while adhering to the substrate, passing from an almost round to a flat shape [16]. The probability distribution of the obtained overall dataset - consisting of 272 measurements - was evaluated by means of a Lilliefors normality test performed on log-transformed cellular dry masses ($\alpha = 0.01$), which demonstrated that it yields a lognormal form (Figure S7B).

The measured probability distribution of cellular dry masses is in agreement with those obtained for unicellular organisms [17]. Although statistically uncorrelated, this outcome also agrees with the 3D lognormality observed for the joint size-metabolism distribution that obtained with these cells in independent experiments (see subsection *Joint measurements of single-cell size and metabolism* in the main text).

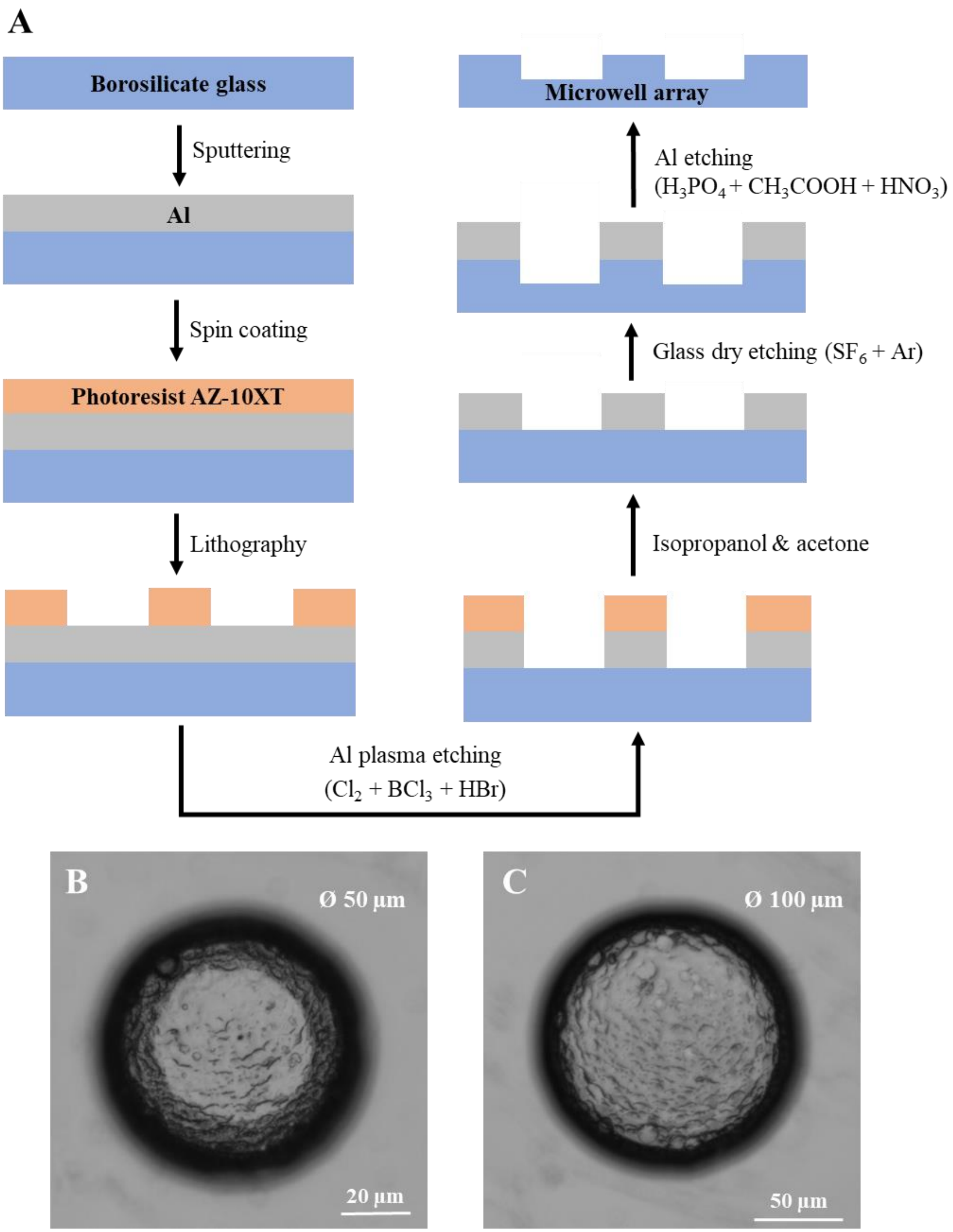


**Figure S1. Microfabrication procedure of a microwell array for performing high-throughput single-cell isolation and readouts. (A)** Based on the desired geometry, a custom aluminum mask for glass dry etching is realized using a standard lithographic process and chemically removed after etching is completed. As a proof of the outcome, brightfield microscopy images of **(B)** a 50 µm-sized (40× objective) and **(C)** a 100 µm-sized (20× objective) microwell immediately after fabrication are shown.

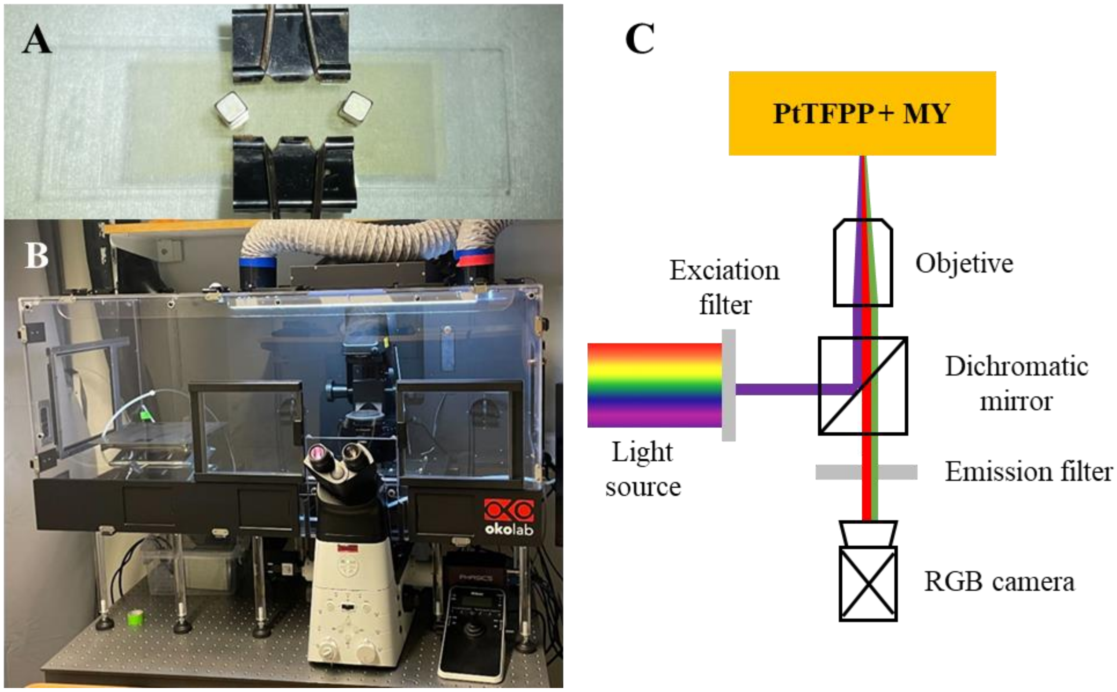


**Figure S2. The experimental set up for performing single cell $O_2$ consumption measurements in the microwell array. (A)** The custom-built glass microwell array**. (B)** The automated microscope surrounded by the transparent incubator. **(C)** Schematic of the optical working principle of the system.

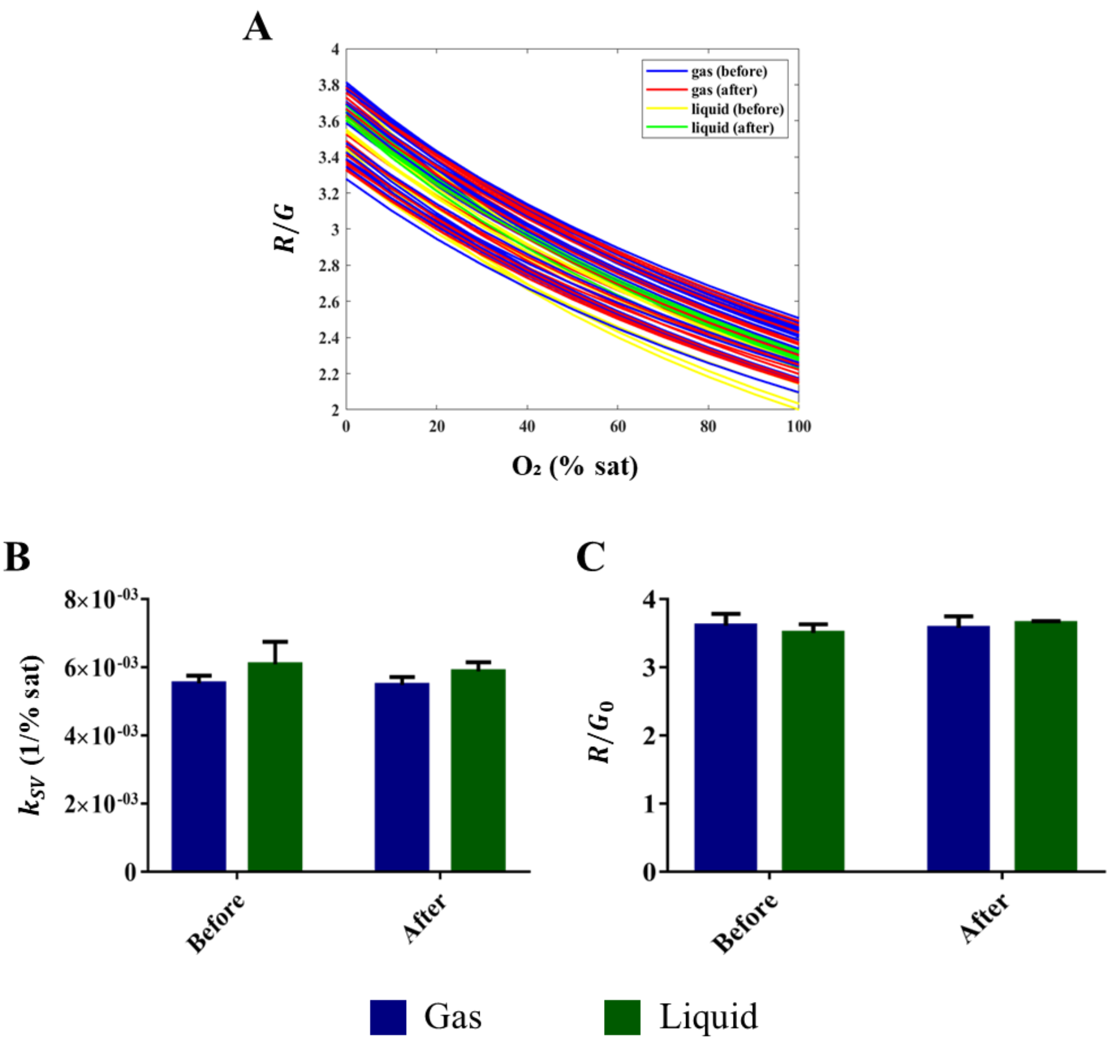


**Figure S3. Assessment of the impact of plasma treatment and calibration phase on optode response. (A)** Calibration curves according to the SV equation, obtained for 20 microwells of the same array before and after plasma treatment filled either with gas or liquid. **(B)** Mean ± standard deviation of the SV constant in the four configurations. **(C)** Mean ± standard deviation of the R/G corresponding to anoxic conditions in the four configurations. No statistically significant differences were observed for either parameter (2-way ANOVA, $p > 0.05$ for both factors of variability).

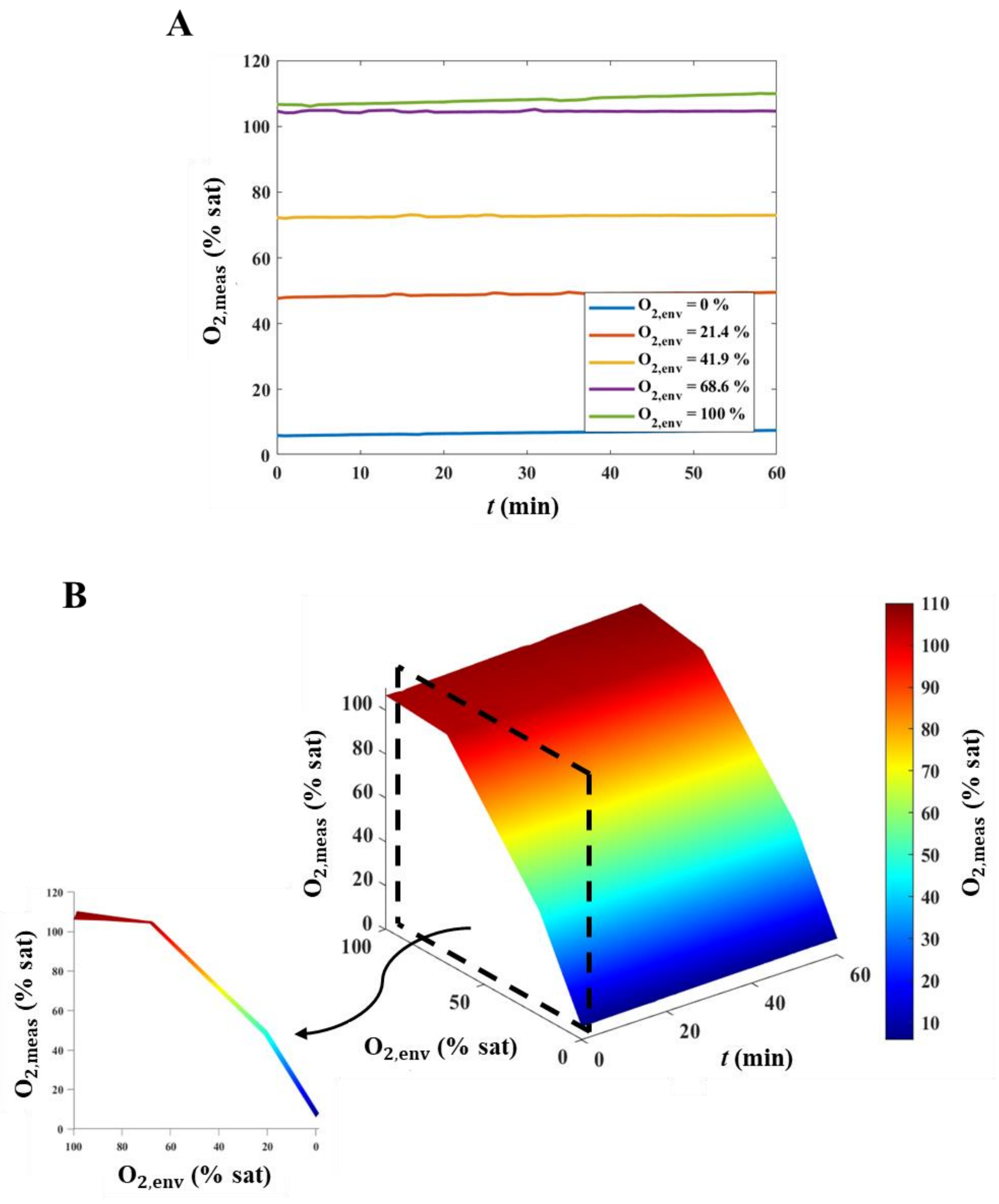


**Figure S4. Characterization of optode bleaching. (A)** Air saturation of $O_2$ measured over time ($O_{2,meas}$), parametrized with respect to the environmental air saturation as set through the gas mixer ($O_{2,env}$). **(B)** Surface plot of the measured air saturation as a function of time and environmental air saturation. Inset: lateral view highlighting the super-linear response of optodes. All graphs refer to air saturation measurements as the average over all 20 microwells considered.

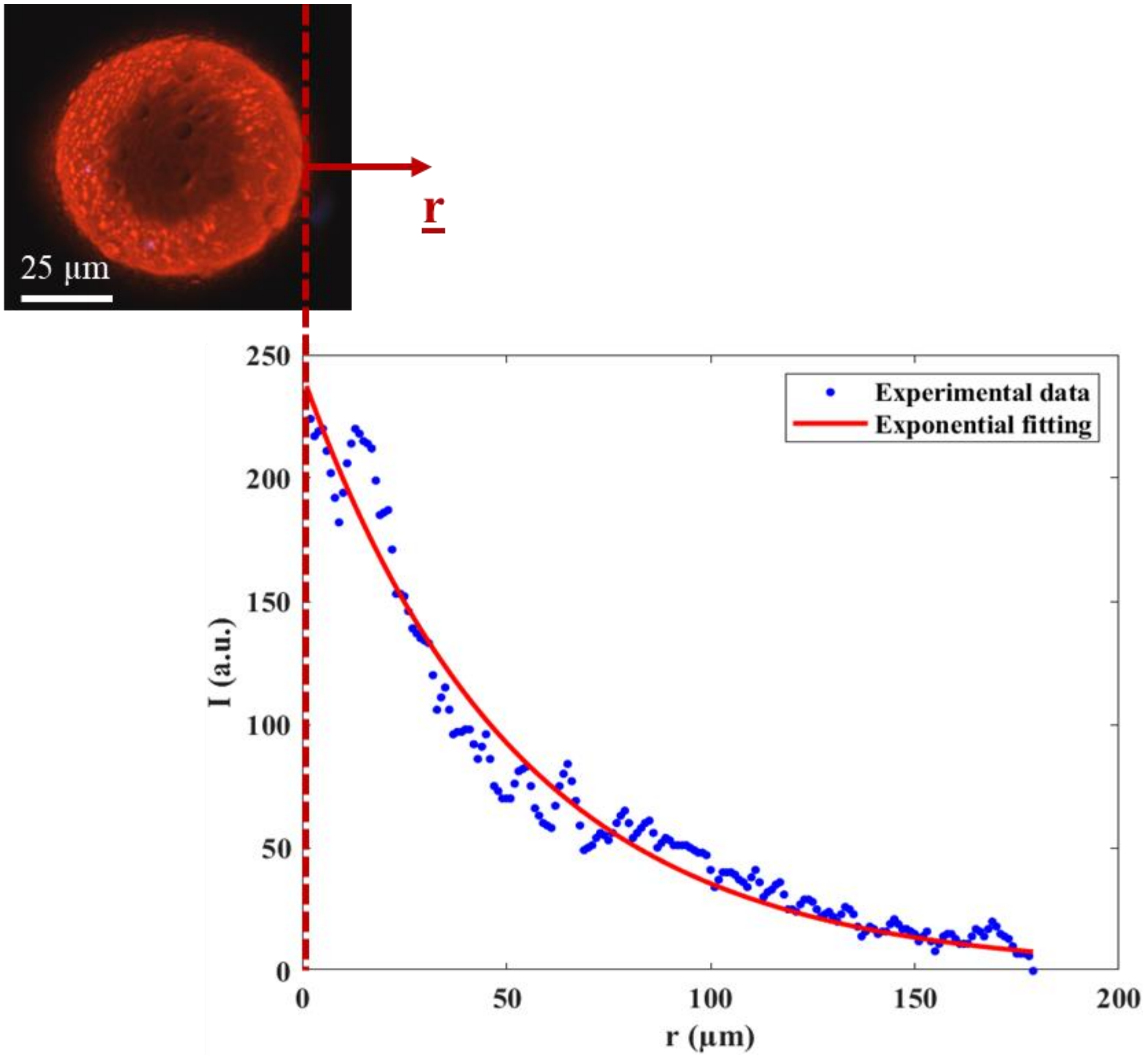

**Figure S5.** An example of the exponential fitting performed on luminescence intensity data for characterizing inter-well optical crosstalk (bottom right). Light intensity is expressed as gray levels of the 8-bit grayscale image corresponding to the red or green channel of the RGB image of a single microwell (top left). The radial axis considered is shown in the small image above the intensity axis.

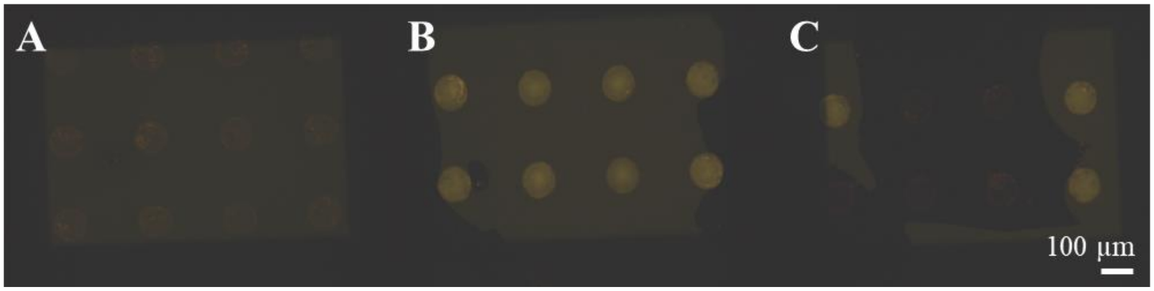


**Figure S6. Heavy mineral oil with Nile Red for the optimization of medium-oil phase separation. (A)** Ideal case of microwells filled with DMEM and overlayed with a homogeneous layer of mineral oil (60 µL each). **(B)** Non-ideal case of microwells filled with mineral oil as medium is displaced (30 µL of DMEM, 60 µL of oil). **(C)** Undesired case of a heterogeneous layer of mineral oil, with some microwells left unsealed (60 µL of DMEM, 30 µL of oil). All images were acquired with a 10× objective and refer to an array with 100 µm-sized microwells.

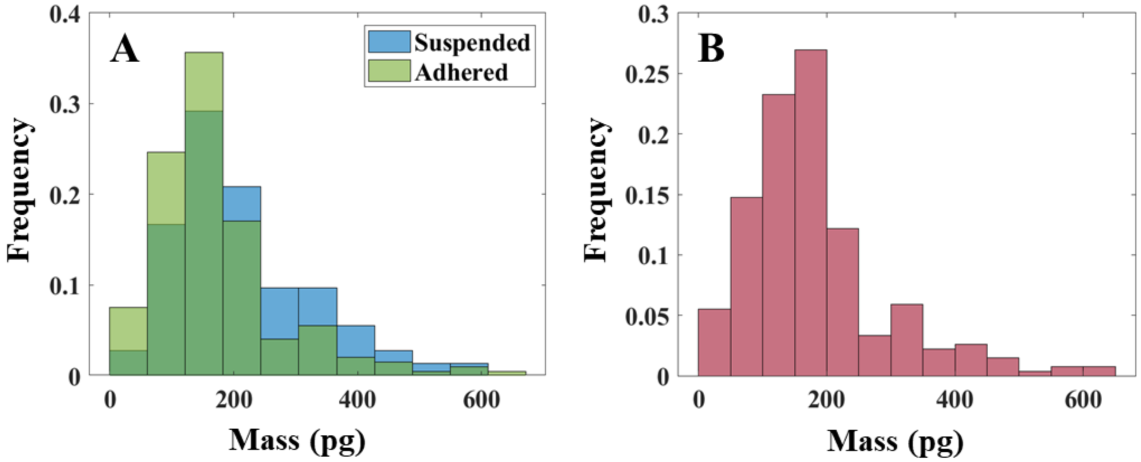


**Figure S7. Dry mass distributions of HepG2 cells measured via a QPI approach. (A)** Mass distributions from images acquired before (suspended, 73 cells) and after (adhered, 199 cells) incubation overnight. The AD test confirmed that their shape cannot be statistically distinguished ($\alpha = 0.01$, $p = 0.0226$). **(B)** Overall experimental mass distribution (272 cells), yielding a lognormal form according to the Lilliefors test ($\alpha = 0.01$, $p = 0.0351$). All histograms are in terms of relative frequencies.

**Table S1.** Technical features of the dry etching process and geometric specifications of the microwell array designed for measuring $O_2$ concentration over time in isolated HepG2 cells.

| Parameter | Numerical value |
|---|---|
| Etching resolution | 1 µm |
| Maximum etching depth | 50 µm |
| Well diameter | 50 or 100 µm |
| Well depth | 20 µm |
| Well volume | 39.3 pL |
| Well-well distance* | Same of diameter |
| Bottom roughness** | 350 ± 150 nm |

**Refers to the distance between the edges of adjacent microwells along the radial direction. ***Refers to the arithmetic average roughness value* [1] *of the bottom surface for an etching depth of 20 µm, expressed as median ± range.*
